# The nature and chromosomal landscape of endogenous retroviruses (ERVs) integrated in the sheep nuclear genome

**DOI:** 10.1101/2022.02.11.480048

**Authors:** Sarbast Ihsan Mustafa, Trude Schwarzacher, JS Heslop-Harrison

**Affiliations:** Department of Genetics and Genome Biology, University of Leicester, Leicester, U.K.; Department of Animal Production, College of Agricultural Engineering Sciences, University of Duhok, Duhok, Kurdistan Region, Iraq

**Keywords:** Chromosomes, Eukaryotic genomes, Endogenous retroviruses, Fluorescent in situ hybridization, Repetitive DNA, Jaagsiekte sheep retrovirus (JSRV), High-throughput sequencing

## Abstract

Endogenous retroviruses (ERVs) represent genomic components of retroviral origin that are found integrated in the genomes of various species of vertebrates. These genomic elements have been widely characterized in model organisms and humans. However, composition and abundances of ERVs have not been categorized fully in all domestic animals. The advent of next generation sequencing technologies, development of bioinformatics tools, availability of genomic databases and molecular cytogenetic techniques have revolutionized the exploration of the genome structure. Here, we investigated the nature, abundance, organization and assembly of ERVs and complete genomes of Jaagsiekte sheep retrovirus (JSRV) from high-throughput sequencing (HTS) data from two Iraqi domestic sheep breeds. We used graph-based read clustering (RepeatExplorer), frequency analysis of short motifs (k-mers), alignment to reference genome assemblies and fluorescent in situ hybridization (FISH). Three classes of ERVs were identified with the total genomic proportions of 0.55% from all analyzed whole genome sequencing raw reads, while FISH to ovine metaphase chromosomes exhibited abundant centromeric to dispersed distribution of these ERVs. Furthermore, the complete genomes of JSRV of two Iraqi sheep breeds were assembled and phylogenetically clustered with the known enJSRV proviruses in sheep worldwide. Characterization of partial and complete sequences of mammalian ERVs is valuable to provide insights into the genome landscape, to help with future genome assemblies and to identify potential sources of disease when ERVs become active.

## 1. Introduction

Mammalian nuclear genomes contain integrated fragments and entire DNA sequences of repetitive genetic elements that have homologous sequences to retroviral genomes known as endogenous retroviruses (ERVs) (Biscotti, Olmo, and Heslop-Harrison 2015; Lander et al. 2001; Mikkelsen et al. 2005; Wicker et al. 2007; Mager and Stoye 2015). ERVs are genetic fossils ascended from the infection and incorporation of ancient retroviruses into the chromosomal DNA of mammalian hosts (Feschotte and Gilbert 2012; Bannert and Kurth 2006). ERVs were revealed in the late 1960s by observing that virological markers could be transmitted following patterns of Mendelian laws (Weiss 2006). Within the repeat element database Repbase, ERV is one of the five superfamilies of LTR retrotransposons which is further subdivided into main five classes; ERV1, ERV2, ERV3, ERV4 and endogenous lentivirus (Kojima 2018). ERVs are largely dispersed in vertebrate genomes in which 8.29% of the entire human genome represents sequences homology to the ERVs fragments (Zhuo and Feschotte 2015; Lander et al. 2001). ERVs are characterized by weak conservation of active sites in the encoded genes and the structure with an internal region including four genes in the order 5’-gag-pro-pol-env-3’. flanked by two Long Terminal Repeats (5’ and 3’LTRs), which act as expression enhancers and/or promoters of cellular genes (Garcia-Montojo et al. 2018; Katoh and Kurata 2013; Bannert and Kurth 2006). In human, most of the ERVs are truncated and silent, although some types of the Human Endogenous Retrovirus (HERV) such as HERV-H, HERV-K, and HERV-W are transcriptionally active (Balestrieri et al. 2015; Grandi and Tramontano 2017). Some retroviruses are associated with diseases: the Jaagsiekte sheep retrovirus (JSRV) is a pathogenic and exogenous retrovirus that have the viral enhancers and promoter in its 5’ and 3’LTRs potentially dynamic in differentiated lung cells and interact with transcription factors specific to the lung that causes Ovine Pulmonary Adenocarcinoma (OPA), an infectious disease in small ruminants (Palmarini and Fan 2001; Arnaud et al. 2007; Sistiaga-Poveda and Jugo 2014). In sheep, the Jaagsiekte sheep retrovirus belongs to the beta related exogenous class II within endogenous groups and therefore is called beta enJSRV (Mager and Stoye 2015; Murcia, Arnaud, and Palmarini 2007). Genomes from many organisms have been sequenced and assembled, but their genome assembly approaches lack efficient method to classify and analyze diversity and genomic distribution of fragmented, degenerated and variable retroviral sequences. Thus, in this study, we aimed to analyze unassembled raw reads from whole genome sequencing of sheep using graph-based read clustering, k-mer frequency analysis and sequence alignments to references, to identify and characterize DNA sequences related to endogenous retroviruses. We aimed to reveal and understand the role of amplification, chromosomal localization and evolution of ERV related DNA repeats and complete genome of enJSRV in the context of genomic and cytogenetic evidence and phylogenetic relationships.

## 2. Materials and Methods

### 2.1 Animal Materials

Blood samples were collected from five individuals of sheep breeds representing two main breeds Karadi and Hamdani from Iraqi Kurdistan region (sample origin in **Table** S1). Total genomic DNA was extracted from whole blood using the Wizard Genomic DNA Purification kit (Promega). Five samples of genomic DNA were sequenced commercially (University of Florida Interdisciplinary Centre for Biotechnology Research) using Illumina NextSeq500 mid-throughput with paired-end 2×150bp cycles, generating 43 to 60 million raw reads (2-3x coverage of the sheep genome) with 5 to 6 Gb total sequence for each DNA sample (Table 5).

### 2.2 Discovery of endogenous retrovirus (ERV)-related repetitive sequences

#### 2.2.1 Graph-based read clustering (RepeatExplorer)

Similarity-based read clustering implemented in RepeatExplorer (Novák, Neumann, and Macas 2010; Novák et al. 2013) was used to identify major repetitive sequences in the whole genome sequence reads. Parameters of RepeatExplorer for clustering included a minimum read overlap length of ≥ 55% of the read length and over 82 bases with 90% of similarity as edges to save the potential error of clustering reads with partial similarity among two unrelated ERV groups. The graphical output clusters with a genome proportion of <0.01% were analyzed to identify retroelement domains and homology hits for the major repeat classes.

#### 2.2.2 k-mer frequency tool (Jellyfish)

As an alternative, reference-free, approach to identify abundant ERV-related sequences, a k-mer analysis of highly abundant sequence motifs k-bases long (k-mers) was carried out: the frequency of all short sequence motifs k-mers in the raw reads were calculated using Jellyfish version 2 (Marçais and Kingsford 2011), with k=22, 32, and 44. The most abundant k-mers in each group were assembled to generate longer contigs representing overlapping k-mers. Several thousand contigs were produced from assembly of short motifs and the top 100 contigs were analyzed.

#### 2.2.3 Database Searching

The consensus sequence of each cluster from RepeatExplorer outcome, and assembled contigs of abundant k-mers, were compared with the Repbase database of repetitive DNA sequences (http://www.girinst.org/repbase/) and NCBI database to investigate and identify similar query sequences to ERV classes of sheep or other animal species.

### 2.3 Design and amplification of FISH probes, chromosome preparation and *in situ* hybridization

Consensus sequences of ERV related contigs were used for designing PCR primer using Geneious software (Kearse et al. 2012) (**Table** 1). PCR amplifications were set up in 25μl total volume reaction mixture containing water (Sigma-Aldrich) (18.4μl), 10x Buffer A (Kapa Biosystems) (2.5μl), 10mM dNTP Mix (1μl), 10μM forward and reverse primers; Sigma-Aldrich (each 0.5μl) and 5U/μL KAPA Taq DNA Polymerase (Kapa Biosystems) (0.1μl) with 80-120ng of genomic DNA. The PCR cycling conditions consisted of 3 min initial denaturation at 95°C, followed by 35 cycles of denaturation (95°C, 0.5min), annealing (Tm-5°C, 0.5 min) and primer extension (72°C, 8min). The final cycle was the 1 min final extension at 72°C followed by indefinite hold time between 4-16 °C. Amplified PCR products were gel electrophoresed (1% w/v agarose) in 1x TAE buffer and purified using an E.Z.N.A. Cycle Pure Kit (Omega) and then labelled with either biotin–16-dUTP or digoxigenin–11-dUTP (Roche Diagnostics) using the BioPrime Array CGH random priming kit (Invitrogen). Probe designations use CL for cluster and ERV for endogenous retroviruses followed by its number. Whole sheep blood was collected from freshly slaughtered commercial sheep (Joseph Morris Butchers Ltd, Leicestershire, UK) in sterile 50ml tubes containing heparin. About 43.5ml of RPMI medium 1640 (GibcoTM, Fisher Scientific, UK); 0.5ml of antibiotic antimycotic solution (10000 lg/ml streptomycin, 10000 U/ml penicillin G, and 25 lg/ml amphotericin B, HyCloneTM, GE Healthcare Life Sciences, Amersham, UK) and 6ml of foetal calf serum were used to make lymphocyte short-term medium. Then, 7ml medium containing either 0.5 or 0.75ml of blood, 10–30mg/ml phytohemagglutinin (PHA; Sigma-Aldrich, UK) was incubated in 5% CO2 incubator at 37°C for 3–5 days. Metaphases were arrested by adding 50–90ml of demecolcine solutions (10 lg/ml; Sigma-Aldrich, UK) and left for further 1.5–2 hours at 37°C. Metaphase chromosome preparations were then made using hypotonic treatment with 0.075M KCl and fixation in absolute methanol: glacial acetic acid 3:1. Fluorescent in situ hybridization (FISH) followed Schwarzacher and Heslop-Harrison (2000). The hybridization mixture contained formamide 50%(v/v), dextran sulphate 20% (w/v), saline sodium citrate (0.3 M NaCl, 0.03 M sodium citrate) 2x SSC, FISH probe 50–100ng, sheared salmon sperm DNA 20μg (Sigma-Aldrich, Dorset, UK), SDS (sodium dodecyl sulphate) 0.3% (w/v) and EDTA 0.12mM. After overnight hybridization at 37oC, low stringency (20% formamide and 0.1xSSC) washes were used enabling probe-target hybrids with more than 70% homology to remain stably hybridized. Biotin-labelled probes were detected with 2.0μg/ml streptavidin conjugated to Alexa594 (Molecular Probes), and digoxigenin probes were detected with 4μg/ml anti-digoxigenin conjugated to FITC (fluorescein isothiocyanate, Roche). Slides were simultaneously counterstained and mounted by applying antifade mixture [6μl DAPI (4’, 6-diamidino-2-phenylindole diluted in McIlvaines buffer pH 7.0; 100μg/ml), 97μl Citifluor antifade mountant solution (Citifluor, Agar Scientific, Stansted, UK) and 97μl ddH2O]. Preparations were analyzed on a Nikon Eclipse N80i fluorescent microscope equipped with a DS-QiMc monochromatic camera (Nikon, Tokyo, Japan) and appropriate filters. Images were false-coloured (red for the probe and cyan for DAPI), overlaid and the contrast adjusted with NIS-Elements BR3.1 software (Nikon) and Photoshop (Adobe) using only cropping and functions affecting the whole image equally.

**Table 1.**
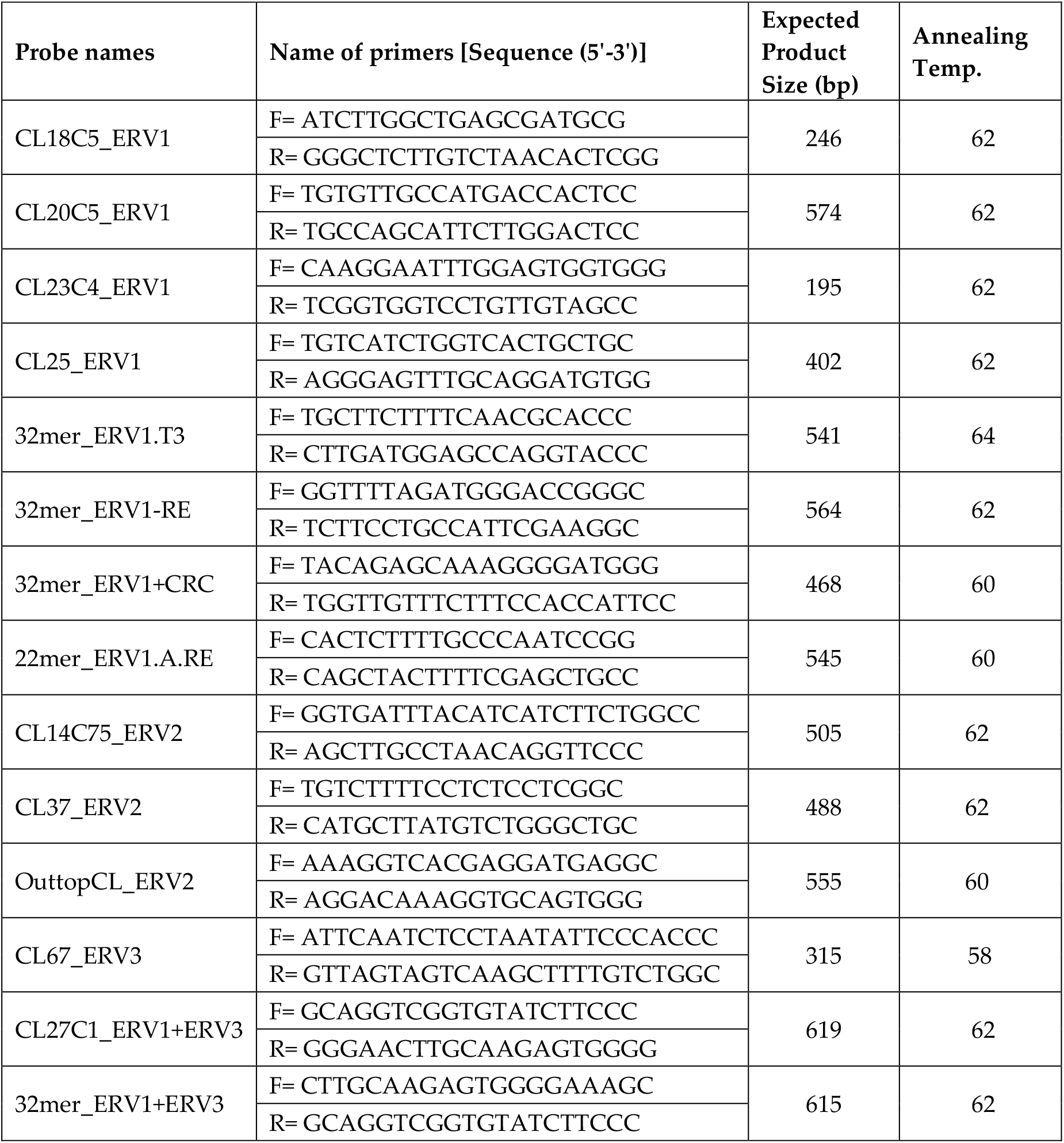
Primer sequences, PCR products, and probe names used for amplification and in situ hybridizations.

### 2.4 Assembly of the complete genome of the endogenous Jaagsiekte sheep retrovirus (enJSRV)

The paired raw reads of the five samples of genomic DNA of Karadi and Hamdani were assembled to the complete genome of the Inner Mongolian Strain of the Endogenous Betaretroviruses Jaagsiekte Sheep Retrovirus (enJSRV; GenBank accession DQ838493) (Wang et al. 2008) to generate consensus sequences for five samples named HamJ1, HamJ2, HamM, KarJ and KarM (Supplementary Materials, Figure S4). The complete genome of enJSRV from Iraqi sheep breeds are available under the GenBank accession numbers of the NCBI MF175067, MF175068, MF175069, MF175070 and MF175071.

### 2.5 Data analysis and phylogenetic relationships

The five complete endogenous betaretroviruses (enJSRV) genomes of Iraqi sheep breeds were aligned with published enJSRV genomes (NCBI) from various sheep breeds worldwide (**Figure** 6). The phylogenetic status was identified by constructing Geneious Tree Builder within the Geneious Prime software. The parameters for the phylogenetic tree were set on Tamura-Nei as a genetic distance model, and Neighbor-Joining as a build method with 500 as a number of replicates. The complete genome of ovine enzootic nasal tumor virus (GU292314) was used as the out-group. Copy numbers, genomic proportions and coverage of ERV sequence (probe fragment) or the complete enJSRV genome were estimated by mapping the whole genome raw reads to the consensus (Table 3, Table 5).

## 3. Results

### 3.1 Identification of endogenous retroviruses related sequences

RepeatExplorer was used to identify and classify LTR retrotransposons by domain sequence homology and order provided by the programme. Several clusters containing ERV sequences were distributed over the RepeatExplorer clusters, each with different genomic proportions (**Table** 2). By comparing the clustered sequences to Repbase databases, all classes of endogenous retroviruses, related to LTR retrotransposons, including ERV1, ERV2 and ERV3 (Bao et al., 2015) were identified. The second approach to identify ERV-related sequences was to use assembled motifs from the most abundant k-mers. Different classes of ERV repeats (ERV1, ERV2 and ERV3) were found by BLAST searches of the top 100 k-mer contigs against repeat databases. Copy numbers and genomic proportion of each probe representing different classes of ERVs related DNA repetitive elements used in this study were estimated following mapping the raw reads against consensus of ERV sequences (**Table** 3). The total genomic proportions of all classes of endogenous retroviruses about 0.55% was estimated based on the genomic abundance of ERV related clusters from the RepeatExplorer outcome.

**Table 2.**
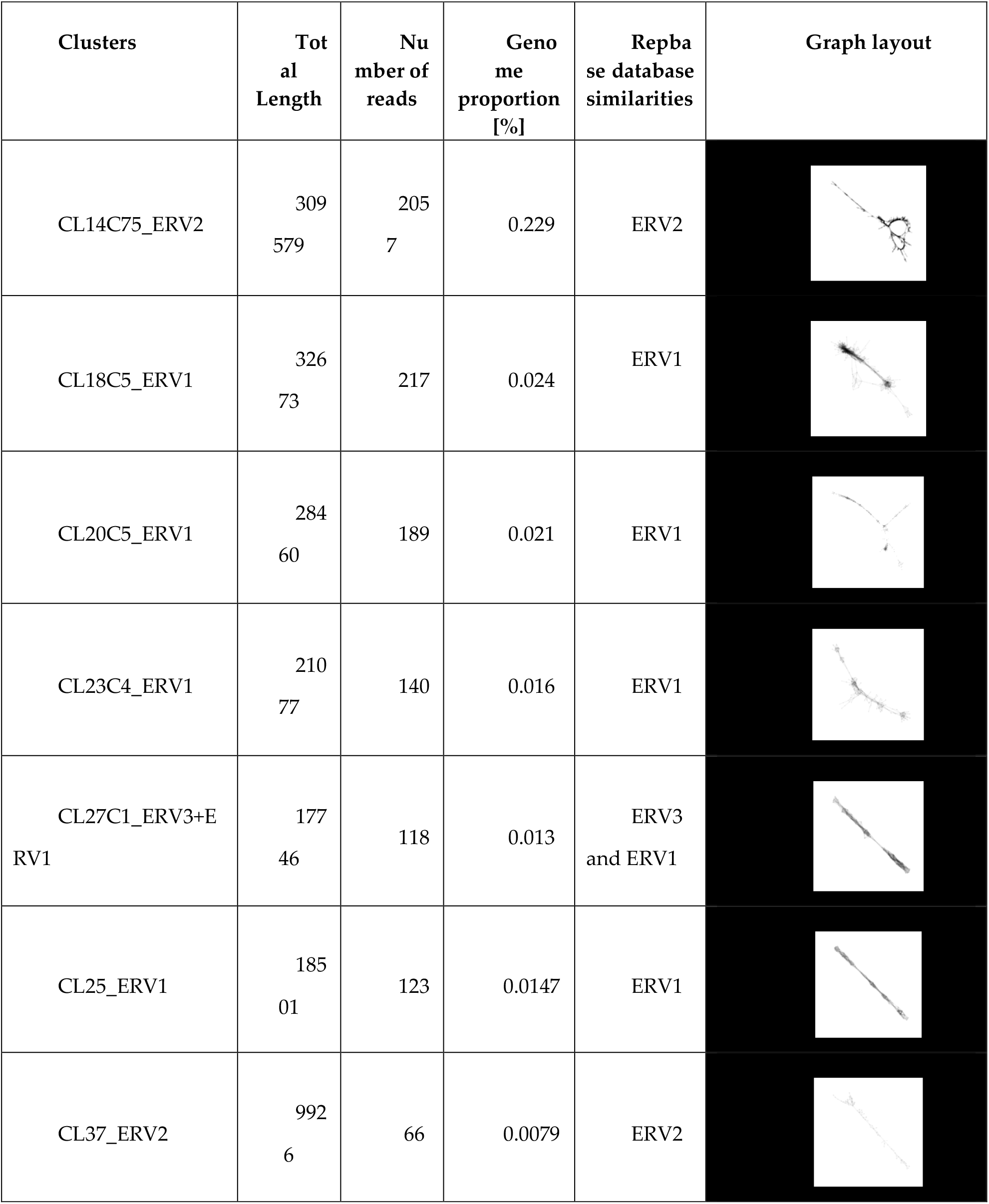
Graph-based clusters with similarity to ERV from RepeatExplorer analysis and Repbase database comparisons.

| Clusters | Total Length | Number of reads | Genome proportion [%] | Repbase database similarities | Graph layout |
| --- | --- | --- | --- | --- | --- |
| CL14C75_ERV2 | 309<br>579 | 205<br>7 | 0.229 | ERV2 |  |
| CL18C5_ERV1 | 326<br>73 | 217 | 0.024 | ERV1 |  |
| CL20C5_ERV1 | 284<br>60 | 189 | 0.021 | ERV1 |  |
| CL23C4_ERV1 | 210<br>77 | 140 | 0.016 | ERV1 |  |
| CL27C1_ERV3+ERV1 | 177<br>46 | 118 | 0.013 | ERV3 and ERV1 |  |
| CL25_ERV1 | 185<br>01 | 123 | 0.0147 | ERV1 |  |
| CL37_ERV2 | 992<br>6 | 66 | 0.0079 | ERV2 |  |

**Table 3.** Copy numbers of various ERV related fragments used for in situ hybridization.

| Probes of endogenous retroviruses ERVs | PCR product bp | HamJ1_Male genome |  |  | KarJ_Female genome |  |  |
| --- | --- | --- | --- | --- | --- | --- | --- |
|  |  | Assembled reads | Copies of probe | Genomic proportion% | Assembled reads | Copies of probe | Genomic proportion% |
| CL15C6_ERV1 | 426 | 3483 | 1226 | 0.0067 | 4024 | 1417 | 0.0066 |
| CL18C5_ERV1 | 246 | 5973 | 3642 | 0.0115 | 7525 | 4588 | 0.0124 |
| CL20C5_ERV1 | 574 | 3124 | 816 | 0.006 | 3676 | 961 | 0.0061 |
| CL23C4_ERV1 | 195 | 2157 | 1659 | 0.0041 | 2000 | 1538 | 0.0033 |
| CL25_ERV1 | 402 | 6900 | 2575 | 0.0133 | 4364 | 1628 | 0.0072 |
| CL39_ERV1 | 270 | 3756 | 2087 | 0.0072 | 3053 | 1696 | 0.005 |
| 44mer_ERV1-T1 | 511 | 5710 | 1676 | 0.011 | 3762 | 1104 | 0.0062 |
| 22mer_ERV1.T2 | 400 | 1720 | 645 | 0.0033 | 2348 | 881 | 0.0039 |
| 32mer_ERV1.T3 | 541 | 1709 | 474 | 0.0033 | 2554 | 708 | 0.0042 |
| 32mer_ERV1-RE | 564 | 3753 | 998 | 0.0072 | 3426 | 911 | 0.0057 |
| 32mer_ERV1+CRC | 468 | 5700 | 1827 | 0.011 | 5760 | 1846 | 0.0095 |
| 22mer_ERV1.A.RE | 545 | 3452 | 950 | 0.0066 | 3008 | 828 | 0.005 |
| CL14C75_ERV2 | 505 | 5294 | 1572 | 0.0102 | 6943 | 2062 | 0.0115 |
| CL37_ERV2 | 488 | 2113 | 649 | 0.0041 | 2338 | 719 | 0.0039 |
| OuttopCL_ERV2 | 555 | 1299 | 351 | 0.0025 | 1177 | 318 | 0.0019 |
| CL67_ERV3 | 315 | 1000 | 476 | 0.0019 | 0 | 0 | 0 |
| CL27C1_ERV1+ERV3 | 619 | 5907 | 1431 | 0.0113 | 5595 | 1356 | 0.0092 |
| 32mer_ERV1+ERV3 | 615 | 5500 | 1341 | 0.0106 | 5300 | 1293 | 0.0087 |

### 3.2 Identification and quantification of ERV repeats in ancestral and *Bos taurus* ERV sequences

Raw reads of sheep were also mapped to concatenated ERVs of 105 ancestral sequences (total length; 145kbp) and 90 sequences of *Bos taurus* (total length; 173kbp) from the Repbase database (Supplementary Materials, Figures S1 & S2). Some 0.02% of the sheep reads (10,518 read pairs) were assembled to the ancestral ERV sequences. BLAST analysis against the Repbase database indicated that these ancestral reads were in type of ERV3 sequences with high (70-88%) similarity. For the concatenated ERVs from *Bos taurus*, 0.45% of the sheep reads (238182 reads) were mapped.

### 3.3 Abundance and genomic organization ERV-related repetitive elements

Amplified PCR products representing selected ERVs were labelled and hybridized to male sheep metaphase chromosomes (2n=54 with three pairs of submetacentric autosomes, 23 autosomal acrocentric pairs, and the X and Y sex chromosomes). Signals varied from dispersed over all or most chromosomes, to localized at centromeres or interstitial positions (**Figure**s 1–5, **Table** 4).

**Figure 1.**
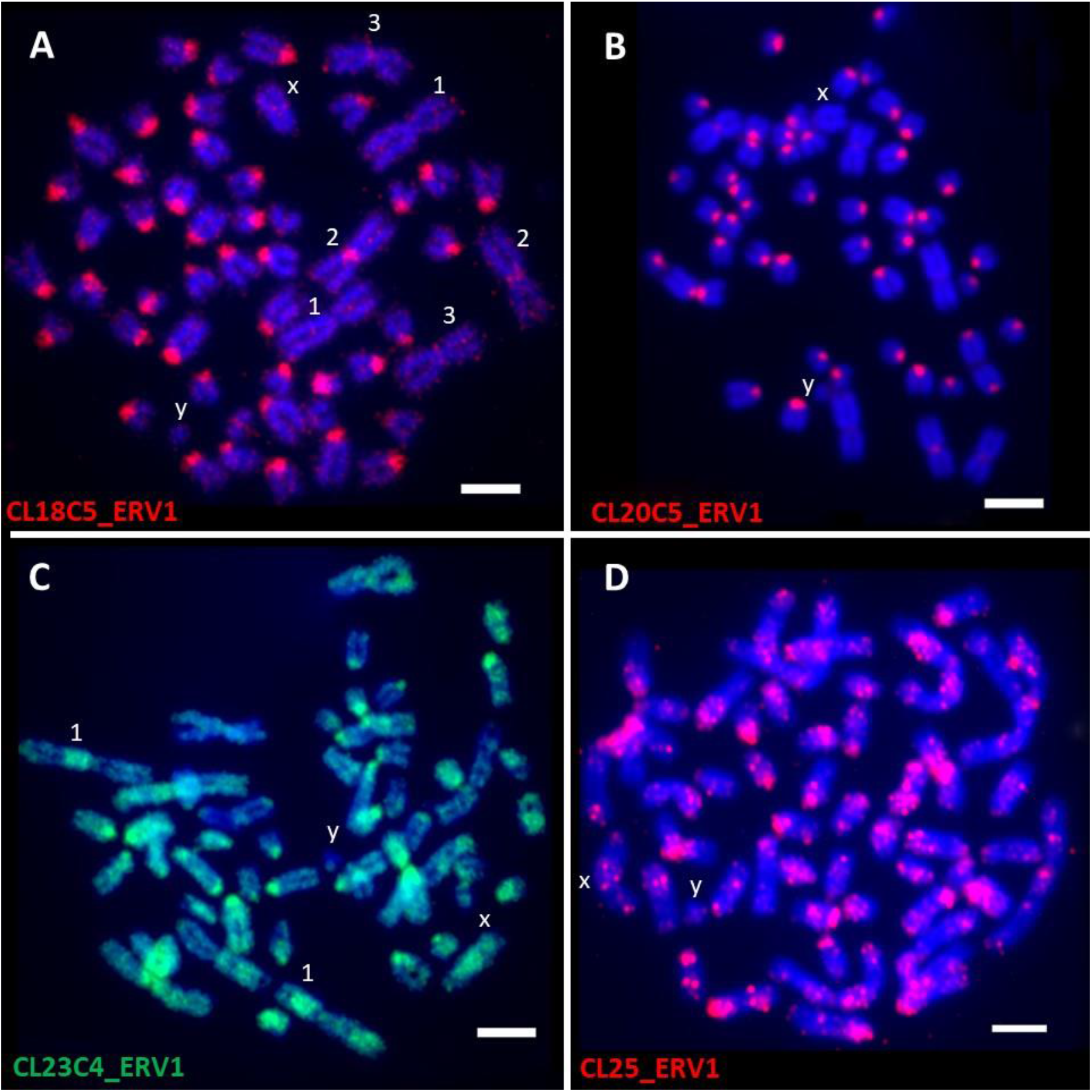
In situ hybridization of ERV1 probes (CL18C5_ERV1, CL20C5_ERV1, CL23C4_ERV1; CL25_ERV1) from Repeat Explorer clusters to the male sheep metaphase chromosomes (2n=54) (fluorescing blue with DAPI). Signals appeared predominantly in centromeric regions.

**Table 4.** Chromosomal characterization of all classes of ERVs

| ERV Probes | Chromosomes | Localization |
| --- | --- | --- |
| CL18C5_ERV1 | All chromosomes except XY and the largest pair of submetacentrics | Centromere |
| CL20C5_ERV1 | All chromosomes except XY and the largest pair of submetacentrics | Centromere |
| CL23C4_ERV1 | All chromosomes | Centromere to disperse |
| CL25_ERV1 | All chromosomes except Y | Centromere and dispersed like dots |
| 32mer_ERV1.T3 | All chromosomes | Centromere to disperse |
| 32mer_ERV1-RE | All chromosomes | Centromere to disperse |
| 32mer_ERV1+CRC | Most chromosomes | Centromere and few telomere |
| 22mer_ERV1.A.RE | All chromosomes | Centromere of all acrocentrics, XY, pair of submetacentrics; Telomere of other 2 pairs of submetacentric |
| CL14C75_ERV2 | All chromosomes | dispersed |
| CL37_ERV2 | All chromosomes | Centromere to disperse |
| OuttopCL_ERV2 | All chromosomes | Centromere to disperse |
| CL67_ERV3 | All chromosomes | Centromere and dispersed like dots |
| CL27C1_ERV1+ERV3 | All chromosomes except XY | Centromere and subtelomere |
| 32mer_ERV1+ERV3 | All chromosomes | Centromere to disperse and few telomere |

#### 3.3.1 ERV1

ERV1 sequences represented in the probe of CL18C5_ERV1 produced strong signals on the centromeres of all 23 autosomal acrocentric chromosome pairs. One pair of submetacentric autosomes had strong centromeric signals, while the other two pairs had weak signals while both X and Y chromosomes had weak centromeric signals (**Figure** 1A). Probe CL20C5_ERV1 showed specific signals at the centromeres of acrocentric autosomes, weaker signals on submetacentric autosomes, and no signals were seen on the largest submetacentrics and sex chromosomes (**Figure** 1B). Signals of probe CL23C4_ERV1 were present on about half the centromeres of acrocentric autosomes, with a more dispersed signals on some but not all chromosomes or arms in the submetacentric chromosomes. There were weak signals on the Y chromosome and stronger signals on the X chromosome (**Figure** 1C). Probe CL25_ERV1 showed variable signals, and slightly dispersed over all chromosomes including the sex chromosomes, with signals close to the centromeres of a few acrocentric chromosomes (**Figure** 1D).

ERV1 sequences from k-mer analysis (22mers and 32mers GT100) also showed centromeric to dispersed patterns on sheep chromosomes: probe 22mer_ERV1.A (**Figure** 2AB) labelled centromeres of all acrocentric autosomes with small dots over the sex chromosomes X and Y, and weak signals or very small dots at centromeric and subtelomeric regions of submetacentrics. Probe 32mer_ERV1 from 32mers GT100 was rather dispersed with a concentration at centromeres, some gaps on submetacentric chromosomes and signals at centromeres of X and Y chromosomes (**Figure** 2C). Probe 32mer_ERV1.T3 from 32mers GT100 showed centromeric and dispersed dots over all chromosomes, mostly rather uniform, including the Y chromosome, while hybridization was more centromeric on the X chromosome (**Figure** 2D).

**Figure 2.**
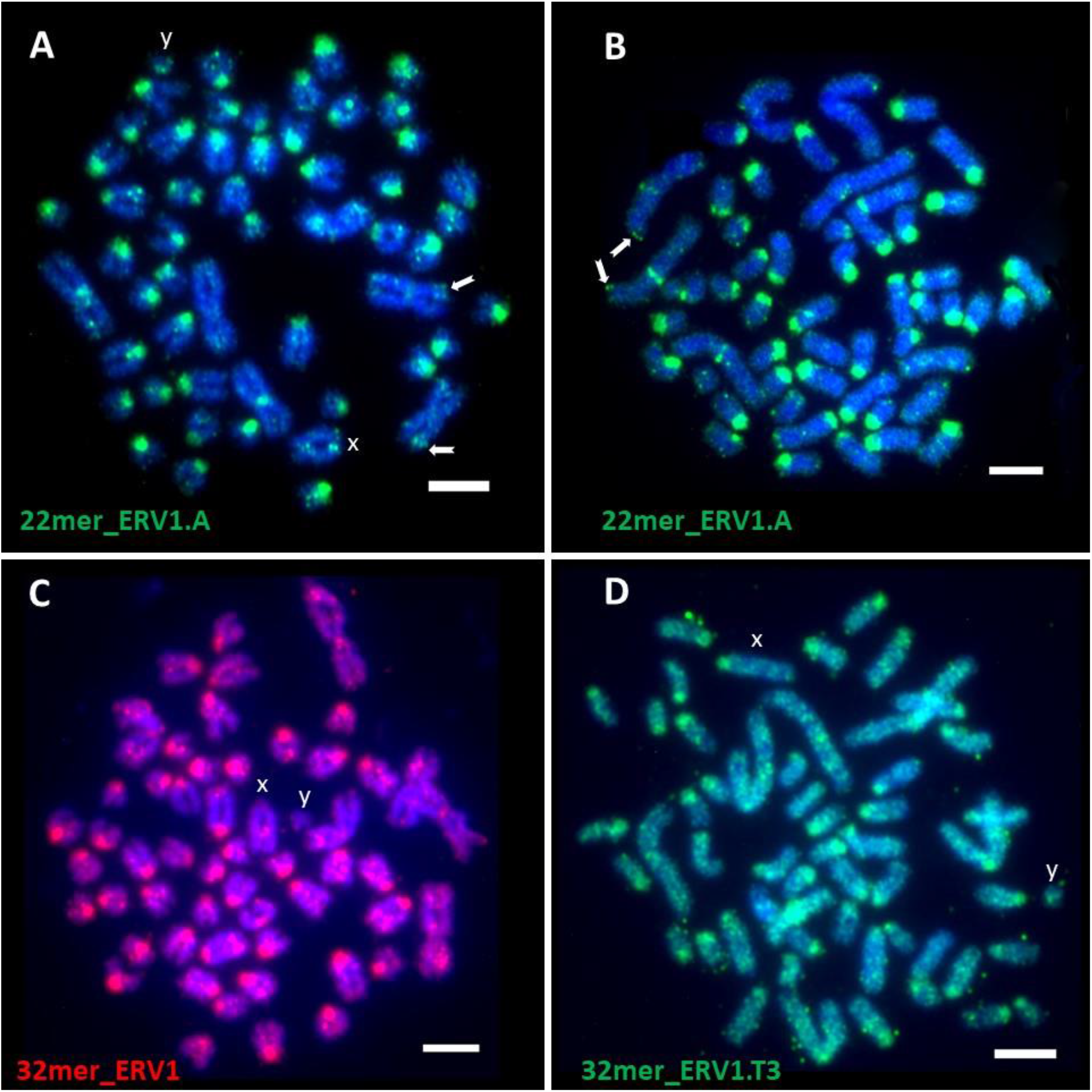
In situ hybridization of ERV1 probes (22 and 32 mer) from kmer analysis to the male sheep metaphase chromosomes (2n=54) (fluorescing blue with DAPI).

#### 3.3.2 ERV2

The chromosomal location of ERV2 showed different patterns to ERV1 (**Figure** 3). Signals of probe CL14C75_ERV2 were broadly dispersed on both autosomes and sex chromosomes, while some DAPI gaps were seen on some chromosomes (**Figure** 3A). The OuttopCL_ERV2 probe showed signals at both centromeric and telomeric domains of acrocentric chromosomes, the submetacentrics had banding-like signals and sex chromosomes had strong signals (**Figure** 3B). Probe CL37_ERV2 showed signals scattered over centromeric regions of some acrocentric and submetacentrics, and slight dot-like signals were seen on all chromosomes including X and Y (**Figure** 3C).

**Figure 3.**
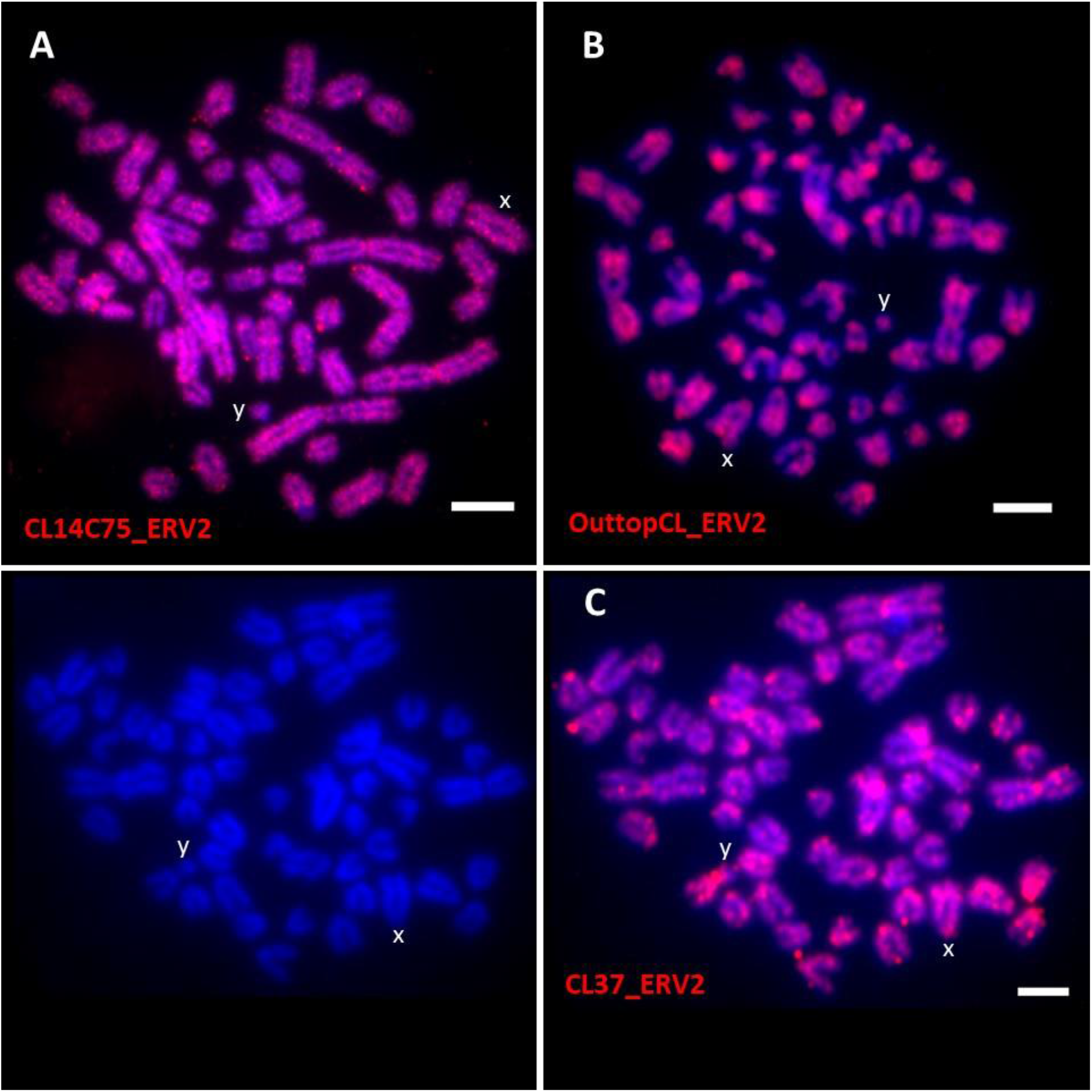
In situ hybridization of ERV2 probes (CL14C75_ERV2, OuttopCL_ERV2; CL37_ERV2) from Repeat Explorer clusters to the male sheep metaphase chromosomes (2n=54) (fluorescing blue with DAPI).

#### 3.3.3 ERV3

ERV3 sequences represented in the probe of CL67_ERV3 formed strong signals at centromeres of all acrocentric chromosomes, and one submetacentric pair, while the other two pairs of submetacentrics and the sex chromosomes had weaker signals (**Figure** 4).

**Figure 4.**
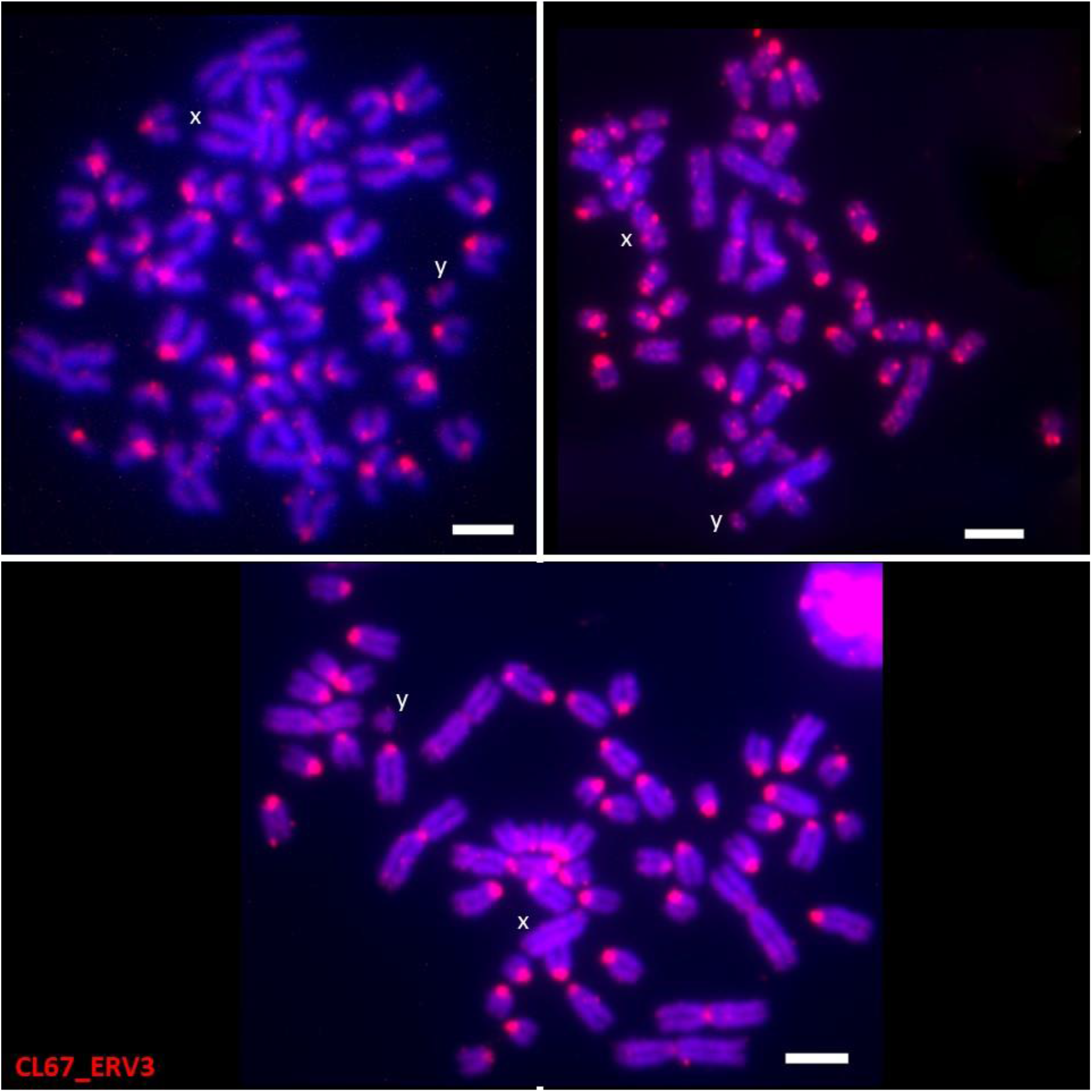
In situ hybridization of ERV3 probes (CL67_ERV3) from Repeat Explorer clusters to the male sheep metaphase chromosomes (2n=54) (fluorescing blue with DAPI). Signals appeared predominantly in centromeric regions.

Consensus sequences of some clusters and k-mer contigs included similarity to both ERV1 and ERV3. The CL27C1_ERV1+ERV3 probe strongly hybridized to the centromeres of all acrocentrics, while centromeric and subtelomeric regions of some submetacentrics had weak signals. Sex chromosomes had no signals (**Figure** 5A). Similarly, probe 32mer_ERV1+ERV3 showed centromeric signals with some bands or broader sites along arms of chromosomes. Telomeric signals on some acrocentrics and submetacentrics were seen. Signals were also incorporated in X and Y chromosomes (**Figure** 5BC).

**Figure 5.**
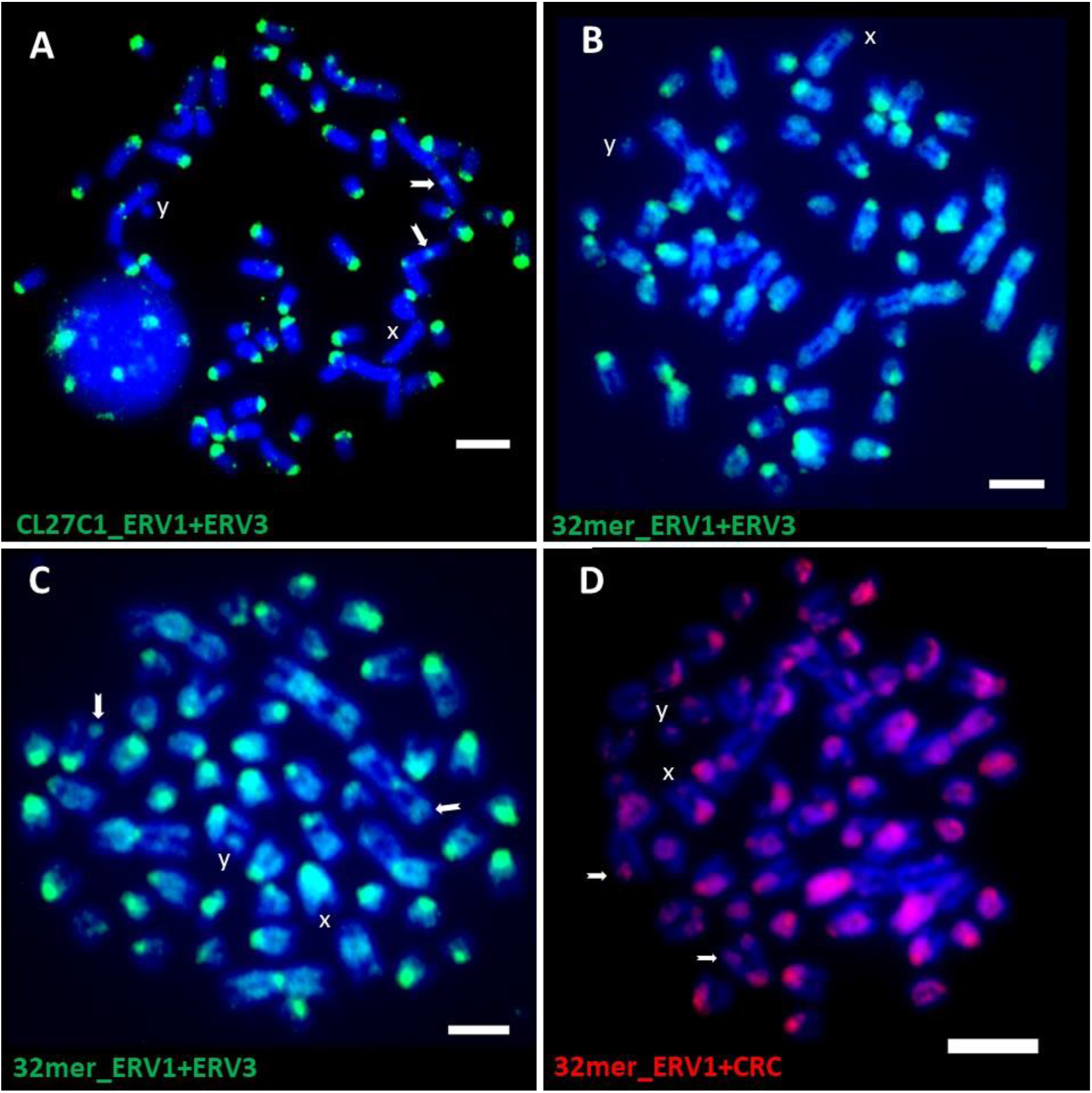
In situ hybridization of combined ERV1+ERV3 probes (CL27C1_ERV1+ERV3; 32mer_ ERV1+ERV3) and ERV1-satellite like sequences probe (32mer_ERV1+CRC) from Repeat Explorer clusters and kmer analysis to the male sheep metaphase chromosomes (2n=54) (fluorescing blue with DAPI).

#### 3.3.4 Combined ERV1 and satellite like sequences

Combined sequences of ERV1 and satellite like repeats was found in the results of both k-mer and RepeatExplorer, suggesting some interspersion of satellite and ERV sequences (Supplementary Materials, Figure S3). Probe 32mer_ERV1+CRC hybridized with the centromeric and the telomeric regions of some acrocentric and submetacentric chromosomes. Some dots were seen on sex chromosomes (**Figure** 5D). Probe CRC (with half of the consensus of 32mer_ERV1 and half CRC) showed 60% similarity to the centromeric repetitive DNA from Cervidae species such as *Muntiacus muntjak vaginalis* (AY064466-AY064469).

### 3.4 The complete genome of the endogenous Jaagsiekte sheep retrovirus (enJSRV)

The complete consensus endogenous betaretroviruses enJSRV genome of three Hamdani and two Karadi sheep breed were assembled. Each was 7941bp long and estimated to be present in 71 to 124 copies (0.0087% to 0.0118% genomic proportion) (**Table** 5). The enJSRV sequences, gene features (predicted proteins, with start and stop codons) and other characteristics such as position, sizes and strand distribution of all 4 protein-coding genes (gag, pro, pol and env genes) and LTR repeats are available in GenBank of the NCBI under accession numbers MF175067, MF175068, MF175069, MF175070 and MF175071. The phylogenetic analysis indicated that the consensus sequences of enJSRV from the large fat-tailed sheep breeds from the Iraqi Kurdistan region was placed with the recognized *Ovis aries* clade including EF680302 and DQ838493 Jaagsiekte sheep retrovirus sampled from geographically different locations (**Figure** 6).

**Table 5.** Copy numbers and genomic proportion of complete genomes of the endogenous betaretroviruses enJSRV integrated in the main sheep breeds of Iraqi Kurdistan region.

| enJSRV Breeds (GenBank ID) | Complete enJSRV genome bp | Assembled reads | Total reads of each genome (Coverage X) | Genomic proportion % | Copies of enJSRV (assembled reads*150/7941) |
| --- | --- | --- | --- | --- | --- |
| HamJ1 (MF175067) | 7941 | 5390 | 52,048,068 (2.6) | 0.0104 | 101.81 |
| HamJ2 (MF175068) | 7941 | 6606 | 56,220,882 (2.81) | 0.0118 | 124.78 |
| HamM (MF175069) | 7941 | 3809 | 43,596,654 (2.18) | 0.0087 | 71.95 |
| KaJ (MF175070) | 7941 | 5386 | 60,605,648 (3.03) | 0.0089 | 101.74 |
| KarM (MF175071) | 7941 | 4846 | 44,933,034 (2.25) | 0.0108 | 91.54 |

**Figure 6.**
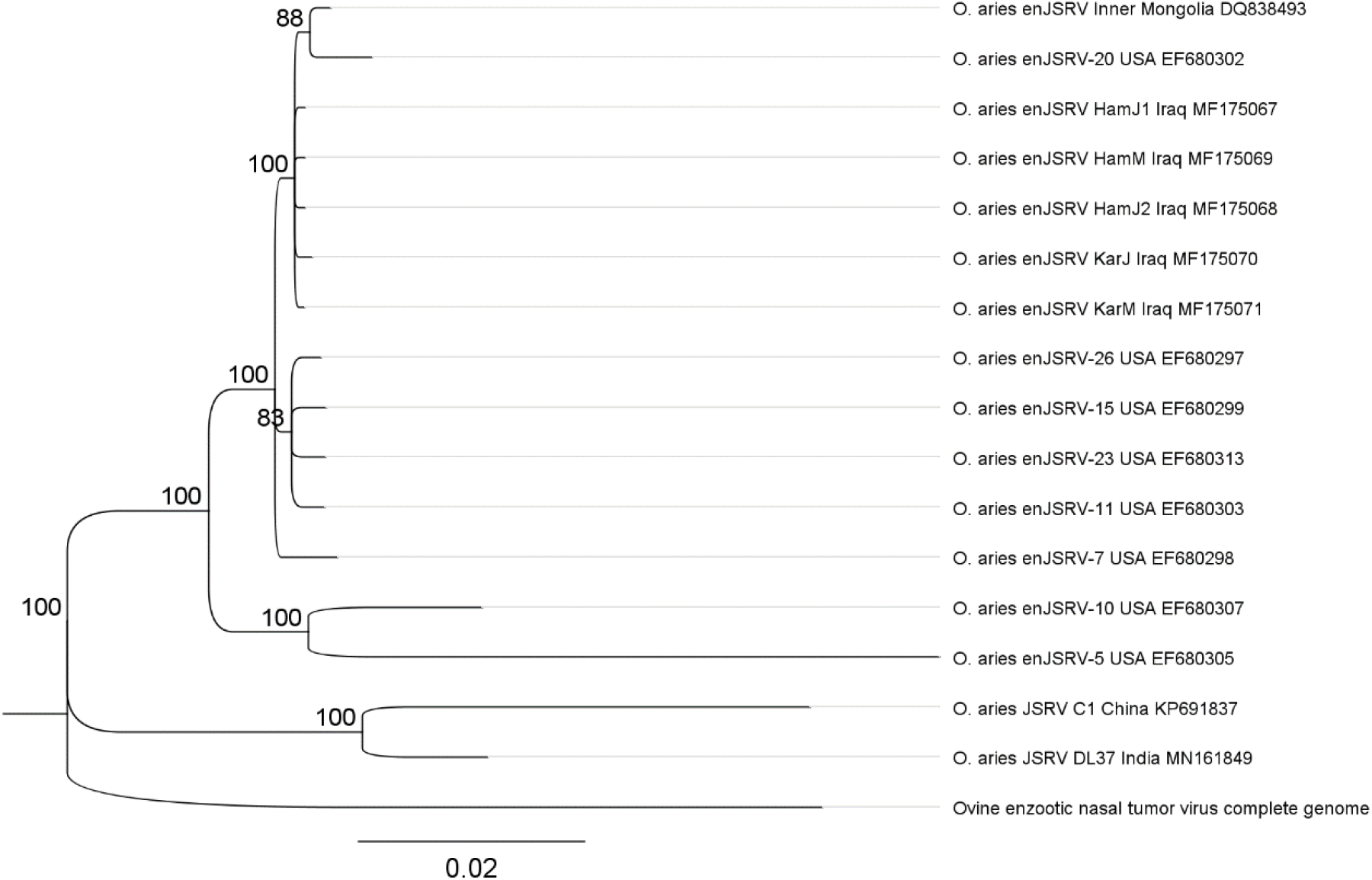
Phylogenetic relationship showing position of complete genomes of endogenous betaretroviruses (enJSRV) of Iraqi sheep breeds (Hamdani; HamJ1, HamJ2, HamM and Karadi; KarM, KarJ) in relation to other strains of enJSRV from other sheep breeds worldwide. Nodes are labelled with consensus support%

## 4. Discussion

### 4.1 Genomic distribution and chromosomal organization of ERVs

Major classes of endogenous retroviruses in the whole genome raw reads of sheep genome were identified using graph-based approaches (RepeatExplorer) (Novák, Neumann, and Macas 2010; Novák et al. 2013) and analysis of abundant k-mers. Fluorescent in situ hybridization (FISH) showed probes from individual ERV sequence families have characteristic but different chromosomal distribution patterns and abundance over the sheep autosomal, acrocentric, metacentric and sex chromosomes. The recognized subfamilies of ERV1, ERV2 and ERV3 showed abundant centromeric to dispersed distribution over sheep chromosomes (**Figure**s 1–5 and **Table** 4). A member of human endogenous retrovirus type-K was found at multiple loci of the pericentromeric locations of several human chromosomes (Zahn et al. 2015), similar to the ERVs at the centromeres of sheep chromosomes. Thus, ERVs are either amplified or accumulate at centromeres, with transposition or retrotransposition and mutation generating to complex family (Prudhomme, Bonnaud, and Mallet 2005). It is notable that the submetacentric autosomes (derived by fusion of acrocentric chromosomes in the ancestral Bovidae karyotype) show contrasting distributions for some ERVs, as has been found for the major centromeric satellite sequences (Chaves et al. 2000). In kangaroo genomes, amplification of endogenous retrovirus occurred in a lineage-specific fashion which is limited to the centromeres of chromosomes (Ferreri et al. 2011). It will be interesting to investigate whether ERVs (in the centromeric and pericentromeric domains depleted in genes) (Lomiento et al. 2008), have any involvement in the karyotypic rearrangement and the evolutionary fusion of the ancestral acrocentric chromosomes, or are carried with the more abundant satellite DNA sequences as has been suggested in bovine chromosome fusions (Escudeiro et al. 2019).

The genomic proportion of ERVs in the sheep genome was estimated as 0.55% in the raw reads from RepeatExplorer, very similar to the proportion aligned to reference bovid ERV sequences from Repbase database (Jurka et al. 2005; Bao, Kojima, and Kohany 2015) (0.02% and 0.45% of sheep raw reads have a high similarity to the ancestral and *Bos taurus* ERV sequences respectively). In three species of Bos; cattle (*Bos taurus*), zebus (*B. taurus* ssp. *indicus*), and yaks (*B. grunniens*), about 30 ERV types were identified and these ERVs were also detected with different percentages in eight species of animals using different types of data (assembly; contigs and reads) (Garcia-Etxebarria and Jugo 2013). In human, approximately 8% of the genome constitutes of retroviral origin sequences, considered a result of continuous infections of the germ line of the host lineage by ancient viruses over millions of years of evolution (Lander et al. 2001; Paces, Pavlícacek, and Paces 2002), much higher than the data here suggests are found in sheep.

The biological significance of ERVs has been argued, but ERVs are now considered to have a variety of beneficial roles in their host genome contributing to genome plasticity (Jern and Coffin 2008; Varela et al. 2009; Kurth and Bannert 2010). In plants, the endogenous pararetrovirus sequences incorporated in the genome, first discovered in banana by *in situ* hybridization (Harper et al. 1999), are now thought to protect the host via RNAi mechanisms (Noreen et al. 2007). Thus, the integrated ERV may have a protective role in reducing retroviral infection in sheep.

### 4.2 Complete genome of endogenous betaretroviruses (enJSRV) and their abundance in Iraqi sheep breeds

Various endogenous retroviruses from different genera have been characterized from a variety of mammalian species (Garcia-Etxebarria and Jugo 2010). Endogenous retroviruses can be categorized through comparison of sequences in a phylogeny (Jern and Coffin 2008). Based on the enJSRV genome, the five assembled Iraqi endogenous betaretroviruses (enJSRV) genomes were grouped together and placed on sister branches with other enJSRV proviruses in sheep within an enJSRV clade (**Figure** 6). enJSRV sequences are thought to have entered the host genome within the last 3 million years, before and during speciation within the genus *Ovis*, and are characterized by a transdominant phenotype able to block late replication steps of related exogenous retroviruses (Arnaud et al. 2007).

About 27 enJSRV proviruses were isolated and characterized in the genomes of different species of genus *Ovis* within Caprinae subfamily including (*Ovis aries, Ovis ammon, Ovis canadensis* and *Ovis dalli*) (Arnaud et al. 2007). Furthermore, copies of enJSRVs and their integration sites in domestic and wild species of the sheep lineage were detected by amplification the env-LTR region by PCR, and 103 enJSRV sequences were produced across 10 individuals and enJSRV integrations were found on 11 of the 28 sheep chromosomes (Sistiaga-Poveda and Jugo 2014) (see FISH results). Similarly, we found that about 294 sequence fragments with different lengths (37bp-7899bp; total length 136kbp) from the nuclear chromosome assemblies of *O. aries Oar_v4.0* databases have high similarity to the genomic sequences of enJSRV. Correspondingly, in the cow genome about 928, 4487, 9698 ERVs related sequences were detected using three different methods, BLAST-based searches, LTR_STRUC and RetroTector respectively (Garcia-Etxebarria and Jugo 2010). The current study is the first work investigating copy numbers 72 to 125 copies and genomic proportions 0.0087% to 0.0118% of complete genome of endogenous betaretroviruses (enJSRV) using whole genome high-throughput sequencing (HTS) data of sheep breeds. The retroviral *pro-pol* sequences of two retroviral families (B/D-type) and (C-type) and the copy numbers (5-100 copies) of B type ERVs in several sheep breeds using Southern blot analysis were analyzed (Nikolai et al. 2003). The ERV sequences identified here fall within the range of diversity previously reported, but form a distinct group (**Figure** 6), suggesting that there is either homogenization of the sequences (including potentially through gene conversion), or loss and replacement with new copies; however, unlike retroelements, it is notable that the copy number of enJSRV is similar across all sheep breeds studied. The results here show that the sheep breeds near the centre of diversity and domestication have similar abundances of enJSRV betaretroviruses and more generally endogenous retroviruses, ERVs to other sheep.

## Supporting information

Supplementary Materials Mustafa et al, 2022

## 5. Availability of data

All the data pertaining to the present study have been included in **Table** and/or **Figure** form in the manuscript and authors are pleased to share analyzed/raw data upon reasonable request.

## Supplementary Materials

The following are available online at xxx

**Table** S1: Sample locations and DNA samples used for **N**ext **G**eneration **S**equencing F; female and M; male.

**Figure** S1: Mapping of whole sequencing of sheep to ancestral sequences of ERVs

**Figure** S2: Mapping of whole sequencing of sheep to *Bos taurus* sequences of ERVs

**Figure** S3: Consensus of CL15C14 (4191bp) of RepeatExplorer including combined ERV1+CRC sequences

**Figure** S4: Assembly of raw reads to reference complete genome of enJSRV DQ838493

## Author Contributions

S.I.M., T.S., and P.H.H. designed and performed the experiments, analysed the data and wrote the manuscript. All authors have read and agreed to the published version of the manuscript.

## Funding

This research was funded by a research studentship from the Kurdistan Region government to S.I.M.

## Acknowledgments

The authors thank the staff of JM Morris, Lutterworth, UK for provision of sheep blood.

## Conflicts of Interest

The authors declare that they have no known conflicts of interest.

## Notes

### Competing Interest Statement

The authors have declared no competing interest.

