## Supplementary Materials Mustafa et al, 2022 for "The nature and chromosomal landscape of endogenous retroviruses (ERVs) integrated in the sheep nuclear genome"

**Table S1.** Sample locations and DNA samples used for **N**ext **G**eneration **S**equencing F; female and M; male.

| Breed[a)] | Sample location [b)] | Sex |
|---|---|---|
| **Hamdani** | **Erbil** | **M** |
| **Hamdani** | **Duhok** | **M** |
| **Hamdani** | **Erbil** | **M** |
| **Karadi** | **Erbil** | **M** |
| **Karadi** | **Duhok** | **F** |

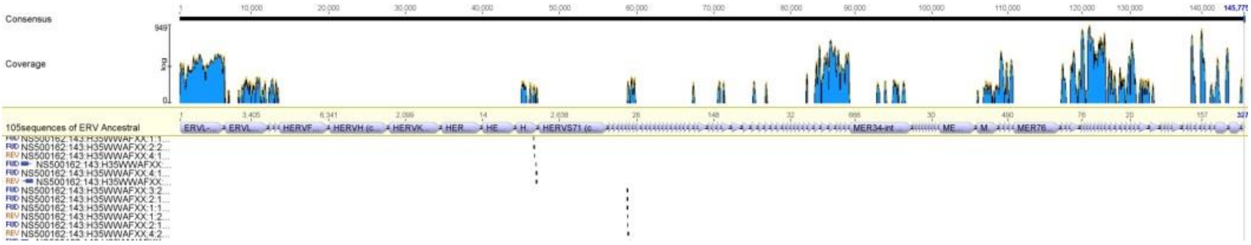


**Figure S1.** Mapping of whole sequencing of sheep to ancestral sequences of ERVs

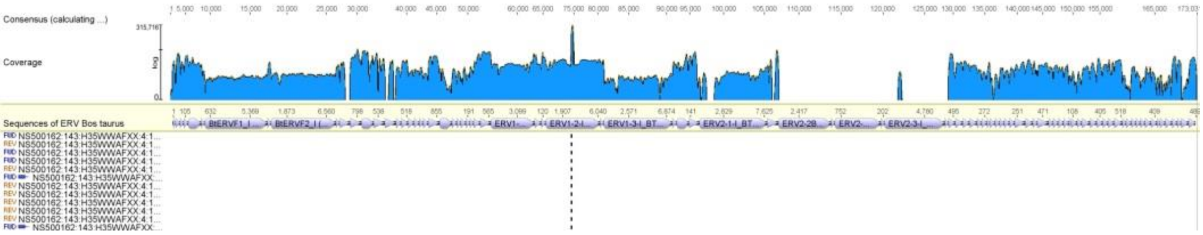


**Figure S2.** Mapping of whole sequencing of sheep to *Bos taurus* sequences of ERVs

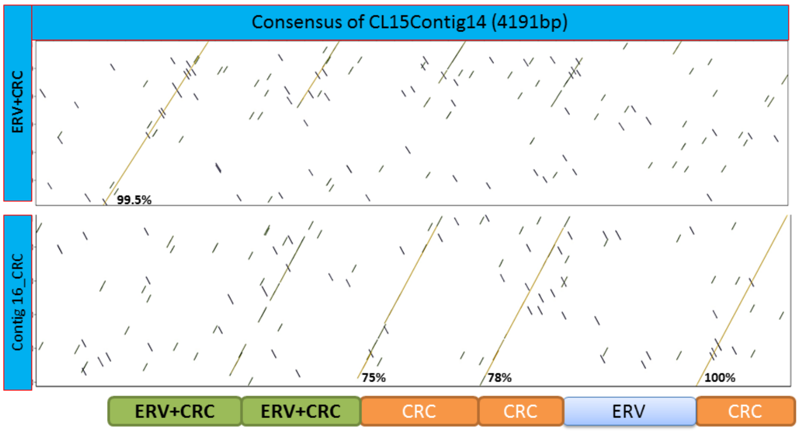


**Figure S3.** Consensus of CL15C14 (4191bp) of RepeatExplorer including combined ERV1+CRC sequences, three copies of 32merC16_Sat_CRC satellite like sequences and

ERVs. Accordingly, the 32merC16_Sat_CRC was named 32mer_ERV1+CRC Figure 5. Furthermore, combined ERV1+CRC sequences and three copies of 32merC16_Sat_CRC satellite like sequences were found in the same consensus of CL15C14 (4191bp) (Figure S3).

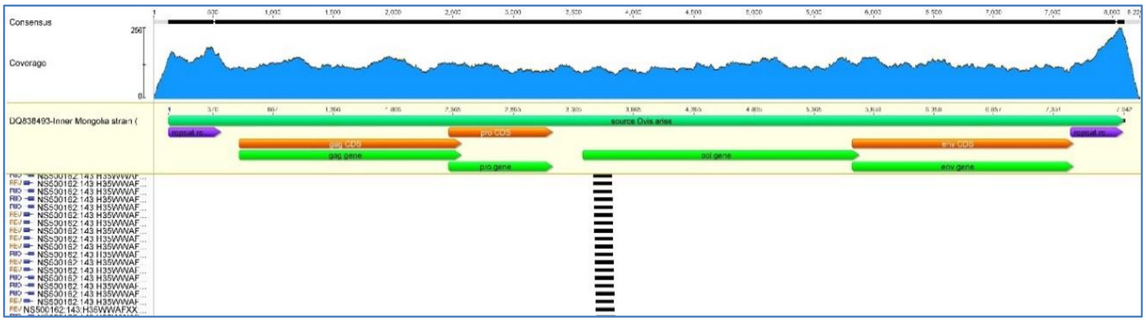


**Figure S4.** Assembly of raw reads (black lines) to reference complete genome of enJSRV DQ838493 showing the 5’ and 3’ LTRs (purple) and the four gene open reading frames, ORFs (brown for coding sequence and genes for green). Reads cover the whole sequence with an average depth of c. 120x (blue, see Table 6.2) and increased depth in the LTRs. Illumina sequencing gave good shotgun coverage (equal forward FWD and reverse REV) reads, and matched left and right paired-end reads to the sequence (shown by symbol after REV/FWD and before read code NS500162…).
